# Conserved Metabolic Regulator ArcA Responds to Oxygen Availability, Iron Limitation, and Cell Envelope Perturbations during Bacteremia

**DOI:** 10.1101/2023.04.18.537286

**Authors:** Aric N. Brown, Mark T. Anderson, Sara N. Smith, Michael A. Bachman, Harry L. T. Mobley

## Abstract

Bacteremia, a systemic infection associated with severe clinical outcomes, is often caused by Gram-negative facultative anaerobes. ArcAB, a two-component regulatory system that represses aerobic respiration, is a key mediator of metabolic adaptation for such bacteria. Using targeted mutational analysis informed by global genetic screens, we identified the *arcA* gene as promoting fitness of *Klebsiella pneumoniae*, *Citrobacter freundii*, and *Serratia marcescens* but not *Escherichia coli* in a murine model of bacteremia. Engineered mutants lacking *arcA* exhibit a dysregulated response to changes in oxygen availability, iron limitation, and membrane perturbations, all of which bacterial cells experience during infection. The genetic response of the *arcA* mutants relative to wild-type strains to the cationic antimicrobial peptide polymyxin B demonstrates an expanded role for ArcA as an activator in response to membrane damage in addition to metabolic adaptation. ArcA function is furthermore linked to electron transport chain activity based on its response to uncoupling of proton motive force by carbonyl cyanide-*m*-chlorophenylhydrazone (CCCP). Differences in lactate and acetate levels as well as lactate dehydrogenase activity between *arcA* mutant and wild-type cells following CCCP treatment establish an ArcA-mediated shift to fermentation independent of oxygen availability. This study highlights the semi-conserved role of ArcA during bacteremia and consolidates infection phenotypes into a comprehensive model based on respiratory activity.

**AUTHOR SUMMARY:** Infections of the bloodstream are life-threatening and can result in sepsis, an overreaction of the host immune system that ultimately damages the body. Gram-negative bacteria are responsible for causing many cases of bloodstream infections, also referred to as bacteremia. The long-term goal of our work is to understand how these bacteria establish and maintain infection during bacteremia. We have previously identified the transcription factor ArcA, which promotes fermentation in bacteria, as a likely contributor to the growth and survival of bacteria in this environment. Here, we study ArcA in the Gram-negative species *Citrobacter freundii*, *Klebsiella pneumoniae,* and *Serratia marcescens.* Our findings aid in determining how these bacteria sense their environment, utilize nutrients, and generate energy while also countering attacks from the host immune system. This information is critical for developing better models of infection to inform future therapeutic development.

## INTRODUCTION

Metabolic flexibility is an established characteristic of opportunistic bacteria and may be a prerequisite for transitioning between non-pathogenic and pathogenic environments. Facultatively anaerobic bacteria are capable of respiration and fermentation and are among the most commonly isolated pathogens from patients with Gram-negative bacteremia (1,2). However, the factors that dictate metabolic shifts during different stages of infection, including colonization and dissemination, are not well understood. *Citrobacter freundii*, *Escherichia coli*, *Klebsiella pneumoniae*, and *Serratia marcescens* cause many community and hospital-acquired cases of bacteremia (3). Bacteremia is often a precursor to sepsis, the single highest cause of in-hospital mortality in the United States (4). *E. coli* and *K. pneumoniae* are the two most frequently isolated pathogens in cases of sepsis while *C. freundii* and *S. marcescens* are emerging bacteremia pathogens of increasing concern (5–8). The long-term goal of this work is to advance our understanding of the metabolic and regulatory pathways employed by these Gram-negative facultative anaerobes within the host bloodstream.

Our group has previously utilized transposon mutant libraries and TnSeq to identify critical fitness genes for *C. freundii*, *E. coli*, *K. pneumoniae*, and *S. marcescens* during bacteremia (9–12). Genes encoding pathways of central carbon metabolism were among the significant fitness genes identified and shared between multiple species. Understanding the regulation of these metabolic pathways is critical for establishing comprehensive models of pathogenesis (13). The TnSeq results were compared between species to identify shared transcriptional regulators of central metabolism that contribute to bacterial fitness. Interestingly, interruption of genes encoding the two-component system ArcAB resulted in a significant loss of fitness for *C. freundii*, *K. pneumoniae*, and *S. marcescens* but not *E. coli*. The response regulator ArcA is a global regulator of metabolism (14) that, together with FNR, IHFA-B, CRP, and Fis, controls the transition between aerobic and anaerobic conditions in the model system *E. coli* (15). Notably, ArcA was the only such regulator found to commonly contribute to bacteremia fitness in *C. freundii*, *K. pneumoniae*, and *S. marcescens*. ArcA has already been shown to be employed by other species including *Haemophilus influenzae* and *Salmonella enterica* in systemic infections (16,17). The most well-studied function of ArcA is repression of aerobic respiration pathways, including the citric acid cycle (18). This regulation is critical for balancing catabolic efficiency (energy production) with fueling anabolism (biomass growth) (19,20). Along with FNR, ArcA controls more than 80% of metabolic flux during fermentation and nitrate-mediated respiration (20). ArcA, and its cognate sensor kinase ArcB, play additional roles in conditions where utilization of available oxygen is suboptimal or potentially detrimental, such as in response to reactive oxygen species (21). Global regulators, including ArcA, are able to integrate multiple stimuli to metabolically reprogram the cell (19,22), and it is likely that several signals in the infection environment may impact ArcA activity. Here, we investigate the role of ArcA in bacteremia by identifying conditions experienced in the mammalian bloodstream that require repression of respiration.

## RESULTS

### Conservation of ArcA

The conservation of ArcA was assessed across the Order Enterobacterales by mapping protein sequences to a predicted structure of ArcA from Alpha Fold (23,24) which is informed by a partially-solved experimental ArcA structure (25). 419 ArcA amino acids sequences (**File S1**) from 418 species across 8 families were identified in total (**Fig. 1A**), with 150 unique ArcA sequences remaining after identical sequences were removed. Conservation analysis based on the ArcA structure and phylogeny of the ArcA sequences calculated an average pairwise distance of 0.07, meaning approximately only 7% of residues differ between any two ArcA sequences. On a scale of 1 to 9, the average conservation level of the 238 residues was 7.7, and more than 75% of residues scored in the “conserved” range of 6 to 9 (**File S2**). The N-terminal receiver domain of ArcA was very well conserved when visualized with pyMOL (26) with the greatest variation observed in alpha helix #2 (**Fig. 1B**). The aspartate residue at the 54^th^ position that is phosphorylated by ArcB in model systems was at the highest level of conservation (27). Between the receiver domain and the subsequent DNA binding domain is a linker domain, which was one of the least conserved regions analyzed. In ArcA, the C-terminal domain is a winged helix-turn-helix (wHTH), which is typical of members of the OmpR family (28). The standard OmpR-like wHTH secondary structure is organized as α_1_-β_1_-α_2_-turn-α_3_-β_2_-β_3_ and is broadly maintained in this model of ArcA (29). The OmpR family of wHTH regulators is characterized by an antiparallel β-sheet upstream of the binding domain which is likely an important determinant of binding specificity (28). The β-sheet of the ArcA model is interspersed with regions of low conservation, suggesting that species-based differences in DNA binding specificity may be reflected in this region. In concordance with the larger sequence comparison, homology of ArcA in representative clinical strains of *C. freundii*, *E. coli*, *K. pneumoniae*, and *S. marcescens* ranged from 93.70% to 99.58% amino acid identity **(Fig. S1)** (30). Evidence identifying ArcA as largely conserved at the amino acid sequence and structural level coupled with the previous genetic screens suggesting *arcA* as supporting pathogenesis prompted the investigation of a shared role during bloodstream infections.

**Fig. 1:**
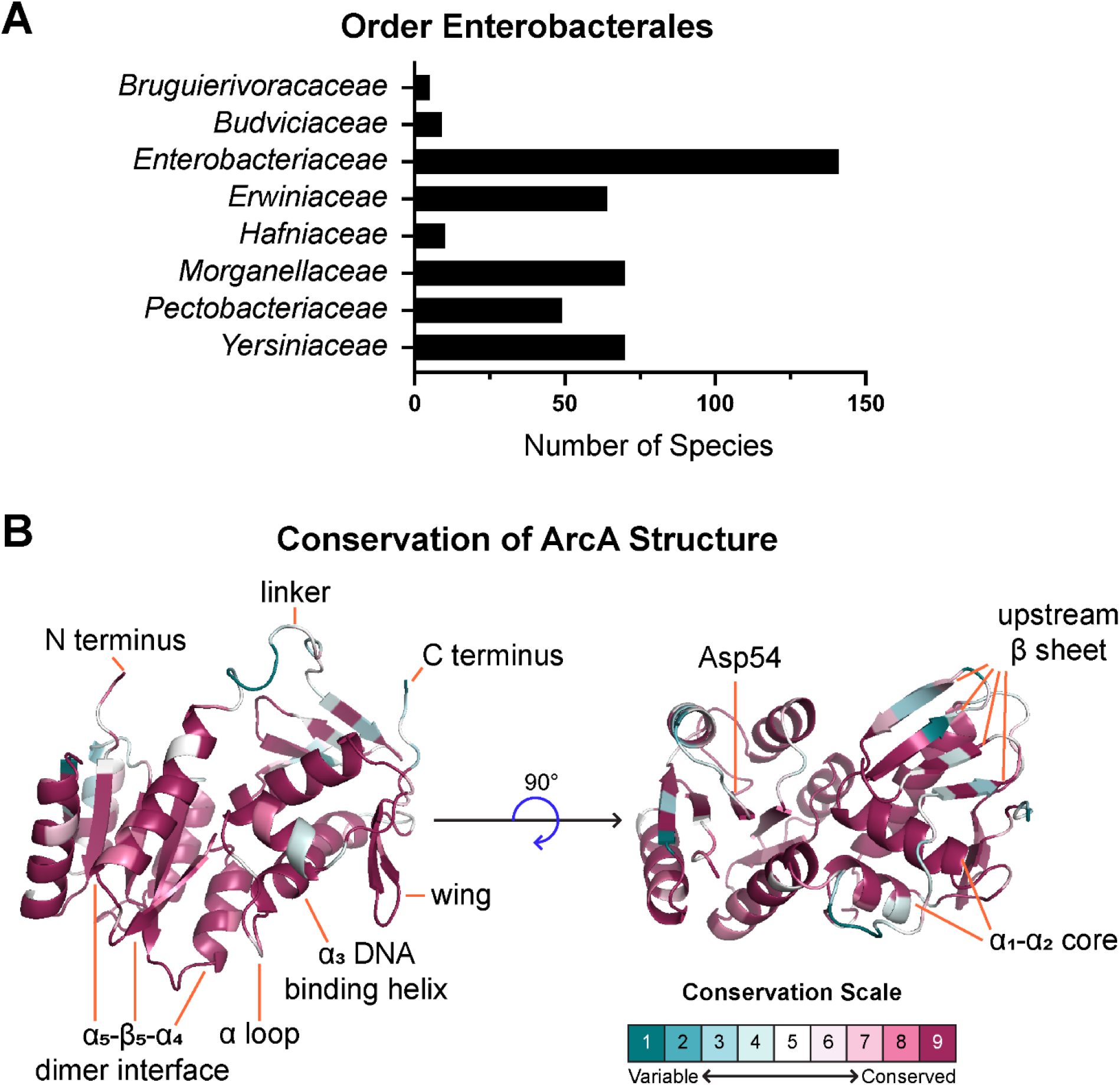
ArcA is structurally conserved across Order Enterobacterales. **(A)** 419 ArcA amino acid sequences of 418 species across 8 families in Order Enterobacterales were identified with BVR-RC and aligned (**File S1)**. **(B)** The multi-sequence sequence alignment was mapped onto a structure of ArcA with Consurf and visualized with pyMOL. The average grade of conservation for 238 residues on a scale of 1 to 9 was 7.7. The regions with the greatest variation in conservation are the linker domain and the upstream β sheet of the DNA binding domain. ArcB activates ArcA via phosphorylation of Asp^54^ which is highly conserved among the species examined in addition to the DNA binding helix and structures supporting it. Conservation of individual residues are available in **File S2**.

### Contribution of *arcA* to fitness in murine bacteremia model

Competition experiments between wild-type and *arcA* mutants **(Table 1)** were conducted in a murine bacteremia model to assess the contribution of ArcA to bacterial survival and replication, collectively referred to as fitness. All tested species colonized the liver and spleen 24-hours post inoculation **(Fig. 2A)** and *S. marcescens* additionally achieved high bacterial burdens in the kidneys, consistent with our previous findings (31). A significant *arcA*-dependent fitness defect was observed in the liver and spleen for *C. freundii*, *K. pneumoniae*, and *S. marcescens* (**Fig. 2B)**. The largest fitness defect for *C. freundii* and *K. pneumoniae* was in the liver where *arcA* cells were outcompeted 6.0-fold and 99.4-fold relative to the isogenic wild-type strain, respectively. *S. marcescens arcA* mutant was most outcompeted in the kidneys (33.7-fold), together indicating the magnitude of ArcA’s contribution to fitness in this model is organ-and species-specific. These results validate and confirm our previous TnSeq findings that initially identified the fitness potential of ArcA among a vast pool of transposon mutants (10–12). No significant fitness defect was observed for the *E. coli arcA* mutant in either the spleen or the liver, a notable contrast to the other species. This finding is corroborated by earlier studies in which an *E. coli arcA* transposon mutant was not associated with a significant fitness defect in spleens by TnSeq (9,32). Thus, although the ArcA sequence analysis demonstrates a high level of conservation, the fundamental contribution of *E. coli* ArcA to bacterial fitness during infection differs substantially from that of *S. marcescens*, *C. freundii*, and *K. pneumoniae*. We therefore chose to further characterize the *arcA* mutants of *C. freundii, K. pneumoniae,* and *S. marcescens in vitro* to explore how ArcA contributes to fitness during bacteremia.

**Table 1:**
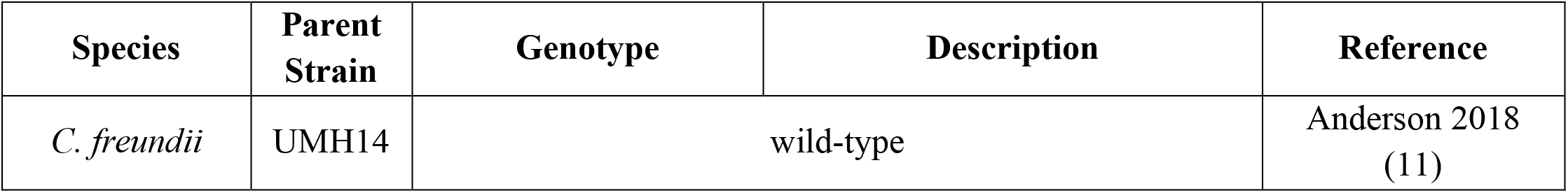
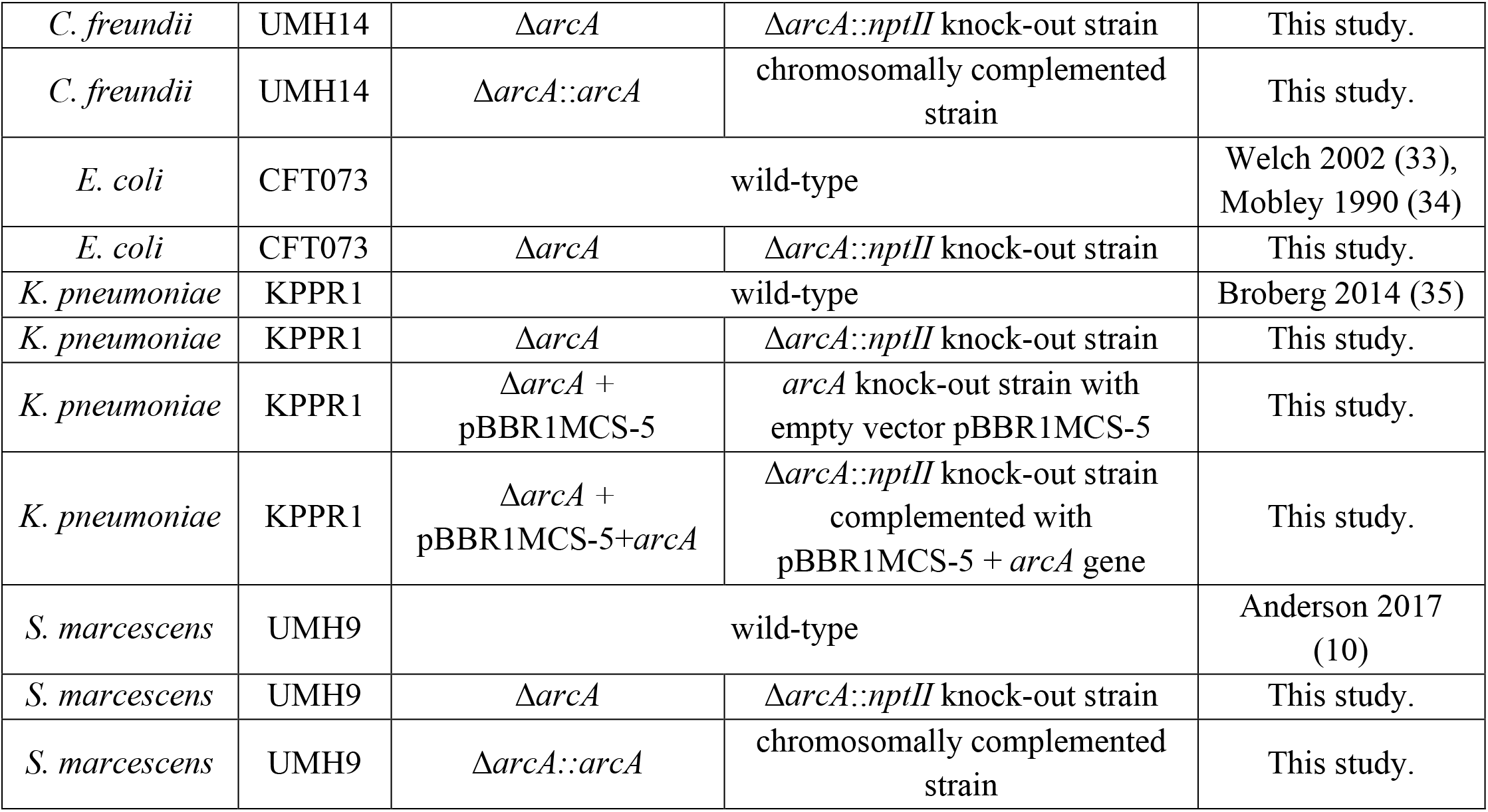
Strains Used in Study

| Species | Parent Strain | Genotype | Description | Reference |
| --- | --- | --- | --- | --- |
| *C. freundii* | UMH14 |  | wild-type | Anderson 2018 (11) |
| *C. freundii* | UMH14 | $\Delta arcA$ | $\Delta arcA::nptII$ knock-out strain | This study. |
| *C. freundii* | UMH14 | $\Delta arcA::arcA$ | chromosomally complemented strain | This study. |
| *E. coli* | CFT073 | wild-type |  | Welch 2002 (33), Mobley 1990 (34) |
| *E. coli* | CFT073 | $\Delta arcA$ | $\Delta arcA::nptII$ knock-out strain | This study. |
| *K. pneumoniae* | KPPR1 | wild-type |  | Broberg 2014 (35) |
| *K. pneumoniae* | KPPR1 | $\Delta arcA$ | $\Delta arcA::nptII$ knock-out strain | This study. |
| *K. pneumoniae* | KPPR1 | $\Delta arcA$ + pBBR1MCS-5 | *arcA* knock-out strain with empty vector pBBR1MCS-5 | This study. |
| *K. pneumoniae* | KPPR1 | $\Delta arcA$ + pBBR1MCS-5+*arcA* | $\Delta arcA::nptII$ knock-out strain complemented with pBBR1MCS-5 + *arcA* gene | This study. |
| *S. marcescens* | UMH9 | wild-type |  | Anderson 2017 (10) |
| *S. marcescens* | UMH9 | $\Delta arcA$ | $\Delta arcA::nptII$ knock-out strain | This study. |
| *S. marcescens* | UMH9 | $\Delta arcA::arcA$ | chromosomally complemented strain | This study. |

**Fig. 2:**
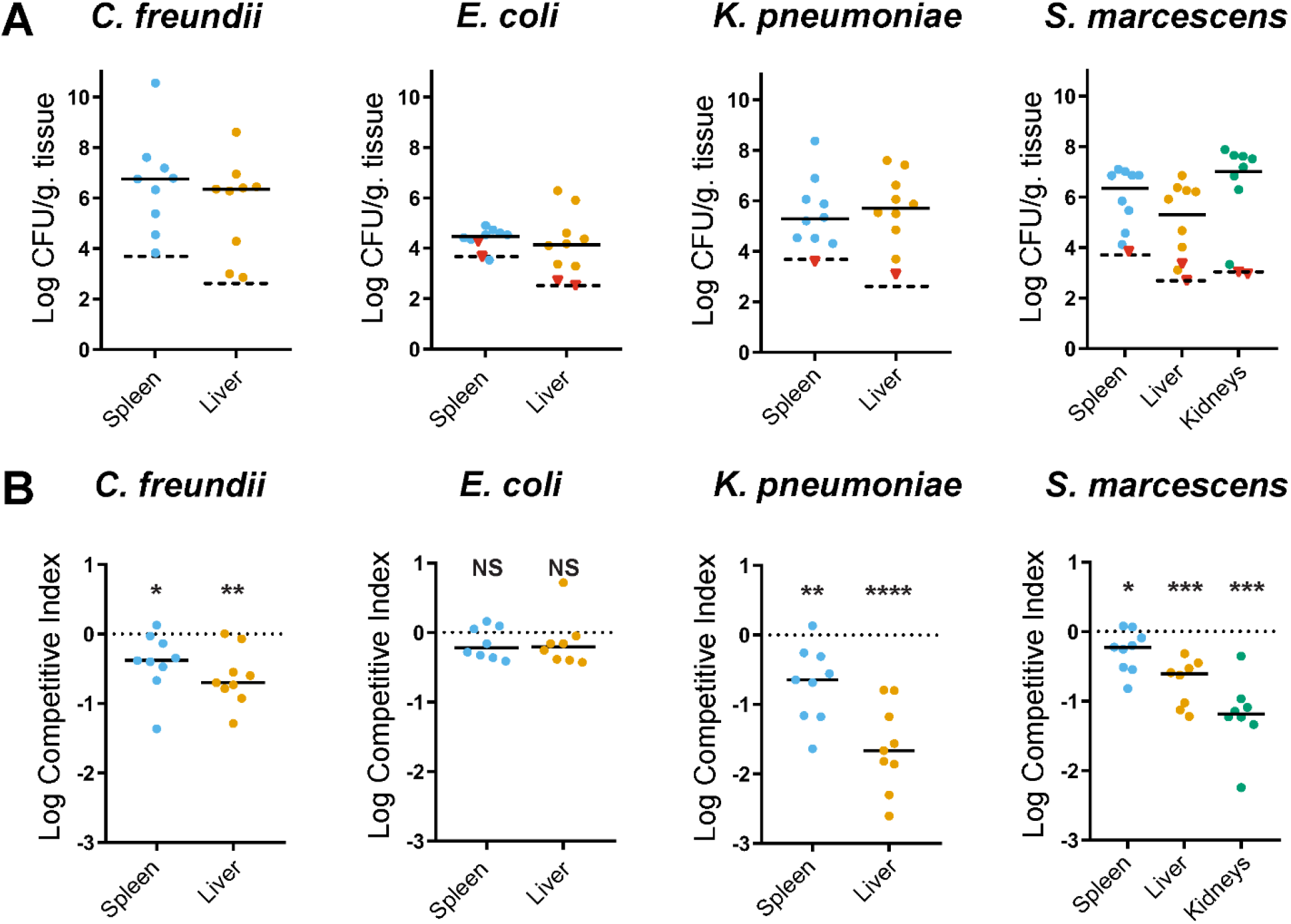
*arcA* encodes a fitness factor in a murine model of bacteremia. Wild-type (WT) and Δ*arcA* mutant strains were cultured to mid-log phase in LB. Cells were washed in PBS and mixed 1:1 to prepare the inoculum for each species at an average target total CFU of 1 x 10^8^ *(C. freundii*), 1 x 10^5^ (*K. pneumoniae*), 1 x 10^7^ (*Serratia marcescens*), and 2 x 10^6^ (*E. coli*). Mice were sacrificed 24 hours post tail vein inoculation, and organs were harvested and plated on LB with and without antibiotics for differential CFU enumeration. **(A)** Total CFU were normalized to tissue weight for all organs. The limit of detection is denoted as a dashed black line, and red triangles are samples not included in calculating competitive indices due to limited CFU recovery. **(B)** Competitive indices (CI) were calculated by dividing the ratio of *arcA* mutant counts to WT counts in the inoculum (input) to that in the organs (output). Dots in the burden and CI graphs represents the organ from one mouse, and median values are presented as solid horizontal lines. Significance of log transformed CI was determined via a one-sample *t*-test with a null hypothetical value of zero, represented as a dotted a line. *p*-values: *≤0.05, **≤0.01, ***≤0.001, NS = not significant

### In vitro growth analysis

The ArcB sensor kinase is classically described as a sensor of anaerobiosis, phosphorylating ArcA under such conditions to optimize growth. *arcA* mutant cells were cultured alongside wild-type and genetically complemented strains to determine ArcA’s influence on bacterial replication in anaerobic conditions across species (**Fig. 3A**). The difference in generation times between wild-type and *arcA* mutant constructs was significant for *C. freundii* (73.5 *vs* 127.1 min.) and *S. marcescens* (113.0 *vs* 173.0 min.) but more modest for *K. pneumoniae* (59.6 *vs* 90.0 min.) **(Table 2**) In recent years, ArcAB has been more precisely described as responsive to a decrease in oxygen consumption (36). To induce a condition in which oxygen utilization is reduced, cells were cultured aerobically overnight and transferred to a strict anaerobic environment before subculturing (**Fig. 3B**). Shifted growth curves from this condition revealed a more substantial delay in the growth of the *K. pneumoniae* and *S. marcescens arcA* mutants compared to the wild-type strains. The *C. freundii*, *K. pneumoniae* and *S. marcescens arcA* mutant strains had 57.5, 22.0, and 72.3 min. longer doubling time relative to the respective wild-type strains after transition from aerobic to anaerobic conditions (**Table 2**). The average doubling time following this transition were very similar to the strict anaerobic condition for *C. freundii* and *K. pneumoniae* strains. These values were considerably longer for *S. marcescens* cells, but wild-type cells continued to grow faster than the *arcA* mutants. Differences in lag time, or the time to reach maximum growth rate, was also calculated (Δ_LT_) as a metric of the cells’ ability to optimize growth performance (**Table 3**). The Δ_LT_ values for *C. freundii* and *K. pneumoniae* were greater in the anaerobic condition, indicating the *arcA* mutant took longer to reach its maximum growth rate relative to the wild-type strain. In contrast, the Δ_LT_ was 29.4 min. longer in the aerobic to anaerobic transition between the *S. marcescens* wild-type and *arcA* mutant strains in comparison to the anaerobic condition.

**Fig. 3:**
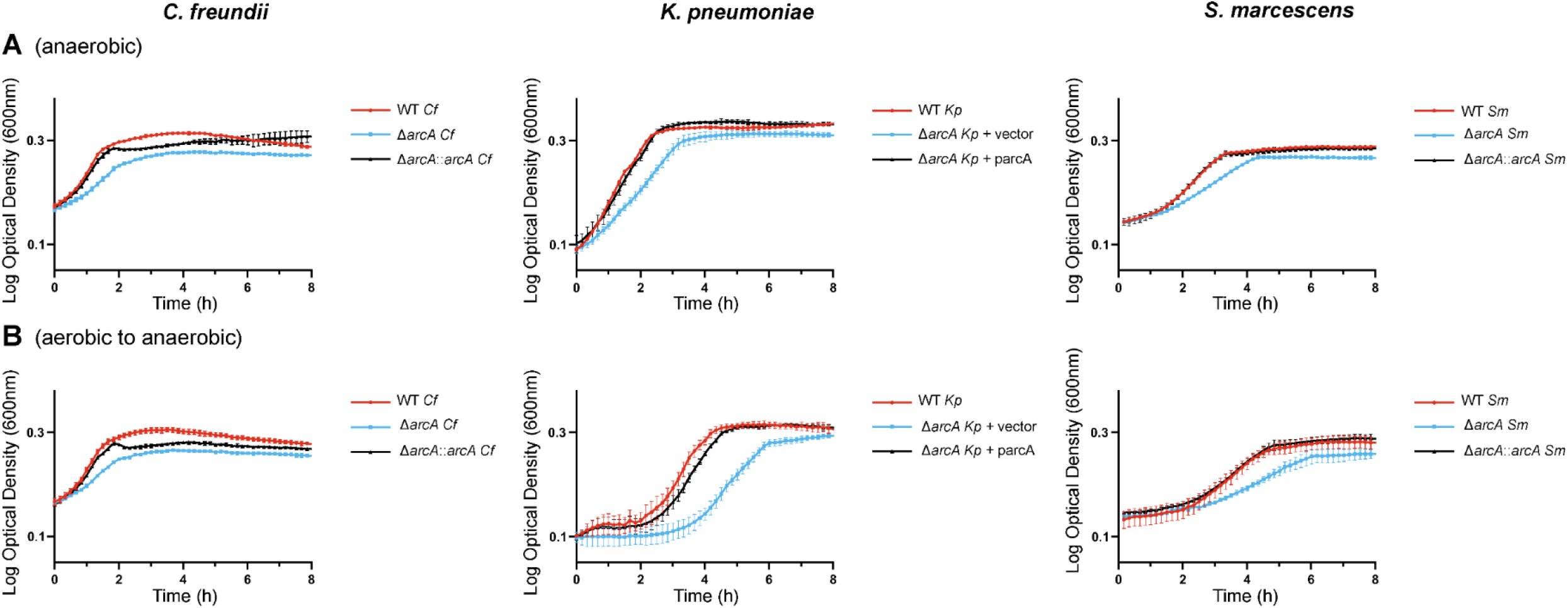
Growth defects of the *K. pneumoniae* and *S. marcescens* Δ*arcA* mutants are more pronounced during the aerobic to anaerobic transition. Strains were grown overnight in LB in (A) anaerobic or (B) aerobic conditions and then normalized based on OD_600_. Fresh LB was inoculated with normalized overnight cultures in an anaerobic chamber. OD_600_ was then measured with a plate reader every 10 minutes. The graphs presented here are representative of three independent experiments. Each strain was grown in triplicate, and the average with standard deviation was plotted over time.

**Table 2:** Doubling times in LB medium (min.)

| Genotype | Anaerobic | Aerobic → Anaerobic |
| --- | --- | --- |
| *C. freundii* |  |  |
| WT | 73.5 ± 5.7 | 72.0 ± 0.7 |
| $\Delta arcA$ | 127.1 ± 7.3 | 129.5 ± 9.8 |
| $\Delta arcA::arcA$ | 80.8 ± 1.2 | 80.3 ± 2.8 |
| *K. pneumoniae* |  |  |
| WT | 59.6 ± 2.9 | 65.1 ± 4.0 |
| $\Delta arcA$ + pBBR1MCS-5 | 90.0 ± 4.6 | 87.0 ± 3.9 |
| $\Delta arcA$ + pBBR1MCS-5+*arcA* | 67.8 ± 5.9 | 73.2 ± 6.8 |
| *S. marcescens* |  |  |
| WT | 113.0 ± 2.7 | 135.6 ± 16.2 |
| $\Delta arcA$ | 173.0 ± 7.7 | 207.9 ± 34.0 |
| $\Delta arcA::arcA$ | 111.0 ± 6.9 | 134.7 ± 4.4 |

**Table 3:**
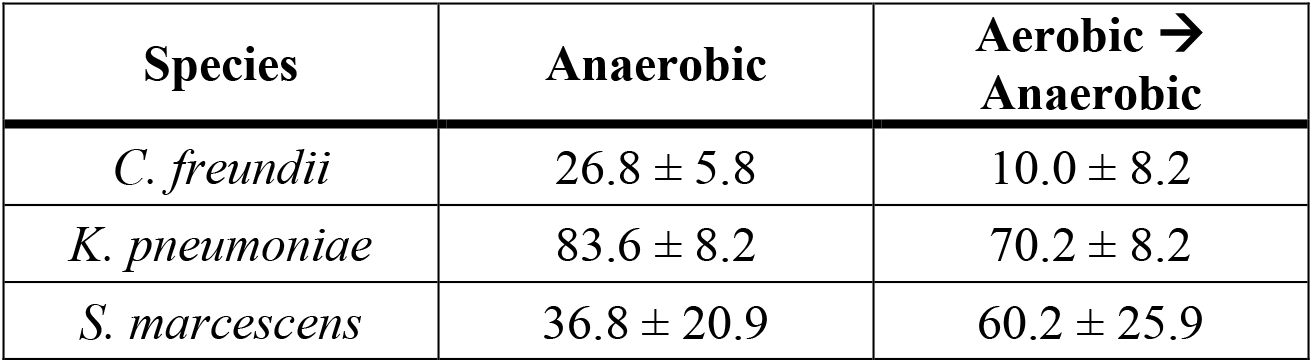
Difference in lag times (wild-type *vs.* Δ*arcA* or *arcA* + eV mutant strains) in LB medium (min.)

Replication of *arcA* mutants was also measured in M9 medium supplemented with glucose and casamino acids to determine if a carbohydrate carbon source alters *arcA*-dependance. The *C. freundii arcA* mutant exhibited a severe growth defect in the presence of glucose for anaerobic culture and aerobic to anaerobic transition culture (**Fig. S2**), for which both phenotypes were more pronounced than in LB medium (**Fig. 3**). Growth defects of the *K. pneumoniae arcA* mutant on the other hand were very similar in glucose-containing medium to those observed in LB. In the presence of glucose, all three *S. marcescens* strains displayed a biphasic growth pattern, with the *arcA* mutant displaying the largest growth defect when bacteria were shifted from aerobic to anaerobic conditions. Overall, the presence of glucose as an available carbon source did not alter the requirement for *arcA* in these three species and indeed exacerbated *arcA*-dependent replication defects for *C. freundii* and *S. marcescens*. The *in vitro* growth kinetics of *arcA* mutants determined here may in part provide a basis for the observed competitive disadvantage of *arcA* mutants during infection, considering that both peptide and monosaccharide carbon sources are expected to be abundant in the host. Furthermore, limited oxygen availability during infection likely plays an important role in how ArcA modulates metabolism of these three species in the bloodstream and tissue environments. However, given the complexity of the infection environment, the potential for ArcA to integrate other relevant signals was also investigated.

### Growth in iron-limited medium

Iron is a critical cofactor for many metabolic enzymes involved in respiration. Enzymes including succinate dehydrogenase and NADH:ubiquinone oxidoreductase require iron-sulfur clusters and are also encoded by operons repressed by ArcA (18,19,37). Free iron levels in the host are low with most iron being bound to hemoglobin and iron-chelating proteins such as ferritin and transferrin (38). During infection, levels of freely available iron drop even further as the host sequesters iron away from the pathogen (39). We hypothesize that ArcA may play a role in metabolic reprogramming in response to iron limitation. Compared to untreated cultures **(Fig. 4A)**, *arcA* mutants grew more slowly than isotypic wild-type strains when cultured aerobically in LB supplemented with the non-utilizable iron chelator 2-2’-dipyridyl **(Fig. 4B)**. Density at stationary phase was considerably lower in the *arcA* mutant cultures in comparison to wild-type cultures. This observation differs from the previous anaerobic experiments where mutant cultures routinely reached the density of the wild-type cells despite any slower growth rates or extended lag periods. Importantly, the phenotype further demonstrates a requirement for ArcA in the presence of oxygen. In all cases, growth kinetics of the three tested species returned to untreated conditions following supplementation of excess iron to dipyridyl-containing cultures (**Fig. 4C)**. The role of ArcA in iron-limited environments is further supported by measuring total growth potential of each species via area under the curve (AUC) in all tested conditions **(Fig. 4D)**.

**Fig. 4:**
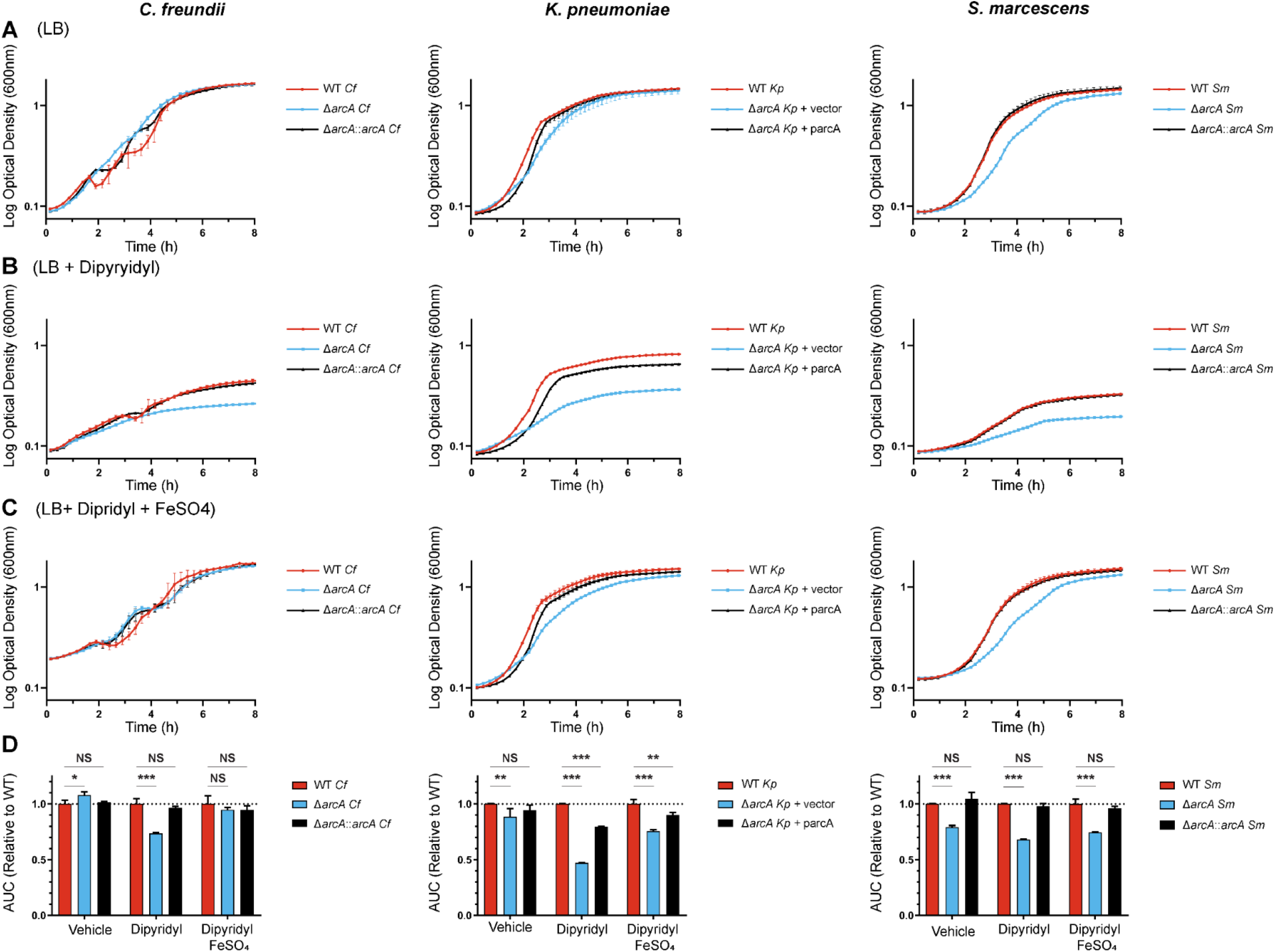
ArcA optimizes growth in an iron-limited medium in aerobic conditions. Overnight cultures incubated aerobically in LB were inoculated into fresh LB containing **(A)** DMSO, **(B)** dipyridyl, or **(C)** dipyridyl supplemented with FeSO_4._ Cultures were incubated at 37°C in aerobic conditions and growth was tracked via OD_600_ by a plate reader every 15 minutes. Growth curves are the average of technical triplicates with standard deviation and are representative of three independent experiments. **(D)** Growth was assessed by calculating area under the curve (AUC) and comparing this value to the AUC of the wild-type in each condition. Bars represent the average of the technical triplicates of the representative growth curves with standard deviation. Significance was determined by comparing the wild-type strain with the mutant and complemented strains with Dunnett’s multiple comparisons test. *p*-values: *≤0.05, **≤0.01, ***≤0.001, NS = not significant

### Sensitivity to killing by human serum

The cell envelope provides the structural barrier necessary to maintain proton motive force generated by the electron transport chain during respiration. Through quinones, the electron transport chain also impacts the kinase activity of ArcB (40,41). ArcA is associated with cell envelope stress in the context of coordination with other envelope regulators, such as σ_E,_ and in direct response to envelope damage (42–44). The bactericidal effects of serum largely target the bacterial envelope (45), and we therefore investigated the role of ArcA in resisting this infection-relevant envelope stress. Viability of wild-type and *arcA* mutants was quantified in the presence of pooled human serum as well as heat-inactivated serum. The *C. freundii arcA* mutant was 37.7-fold more susceptible to killing by intact serum relative to the wild-type strain (**Fig. 5A**), a phenotype partially complemented in the Δ*arcA*::*arcA* strain. In contrast, all three *C. freundii* strains exhibited growth in culture with heat-inactivated serum, but the *arcA* mutant did not grow as robustly as the wild-type and *arcA* complemented strains. None of the *K. pneumoniae* strains exhibited reduced viability when cultured with 90% human serum, demonstrating a high level of serum resistance for this strain (**Fig. 5B**). Interestingly, the wild-type and complemented *arcA* strain *K. pneumoniae* grew to similar levels in heat-inactivated serum while the *arcA* mutant showed a significantly reduced ability to replicate in the serum environment. Serum-mediated cell death was also observed in the *S. marcescens* strains in 40% serum where the *arcA* mutant experienced a 16.7 times more killing relative to the wild-type strain (**Fig. 5C**). The *S. marcescens* strains cultured in the heat-inactivated serum experienced net growth rather than killing to similar levels as the *C. freundii* strains except no statistical difference between wild-type and *arcA* mutant strains was observed. Disparities in growth between mutant and wild-type strains in heat-inactivated serum suggests that nutrient limitation or another growth condition inherent to serum likely contributes to these results. Nevertheless, ArcA contributes to complement resistance for *C. freundii* and *S. marcescens*, demonstrating the link of this response regulator to membrane integrity for these species.

**Fig. 5:**
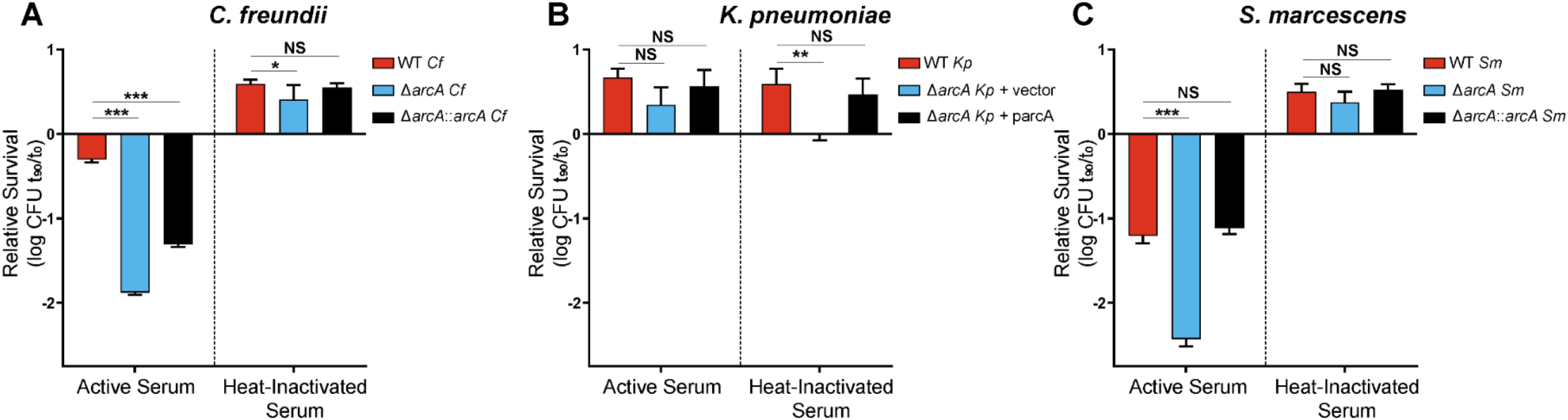
ArcA is required for serum resistance of *C. freundii* and *S. marcescens*. Overnight cultures incubated in LB medium were sub-cultured into LB medium and incubated aerobically until mid-log phase. Cells were normalized and resuspended in active and heat-inactivated human serum to a final concentration of approximately 2 x 10^8^ CFU/mL. Cultures were then incubated at 37°C for 90 minutes with sampling before and after incubation for CFU enumeration. Each species was treated with an empirically determined concentration of human serum at the following final concentrations: **(A)** *C. freundii*: 10%; **(B)** *K. pneumoniae*: 90%; **(C)** *S. marcescens*: 40%. Average values of technical triplicates with standard deviation are presented on each graph and are representative of three independent experiments. Significance was determined by comparing the wild-type strain with the mutant and complemented strains with Dunnett’s multiple comparisons test. *p*-values: *≤0.05, **≤0.01, ***≤0.001, NS = not significant

### Response to polymyxin B

The host innate immune response includes cationic antimicrobial peptides (CAMPs), such as cathelicidin LL-37, which permeabilize bacterial cell membranes (46). Polymyxin B (PMB) is a model CAMP and was used to test whether ArcA also plays a role in the response to CAMP-mediated cell membrane damage (47,48). PMB treatment of mid-exponential phase cells demonstrated that *arcA* mutants of all three species were significantly more susceptible to killing than their isogenic wild-type strain and complemented mutants **(Fig. 6A)**. Survival rates were 44-, 138-, and 76-fold higher in the wild-type strains relative to the a*rcA* mutant constructs of *C. freundii*, *K. pneumoniae*, and *S. marcescens*, respectively. These results are especially notable for *K. pneumoniae* given the lack of *arcA*-dependent serum resistance observed (**Fig. 5B**), thus supporting the conclusion that ArcA also has a role in responding to *K. pneumoniae* membrane perturbation similar to *C. freundii* and *S. marcescens*. Together, these data support previous findings that ArcA regulation of downstream target genes is important for cellular processes that support envelope health. To investigate further, an ArcA-specific genetic response to polymyxin B was interrogated.

**Fig. 6:**
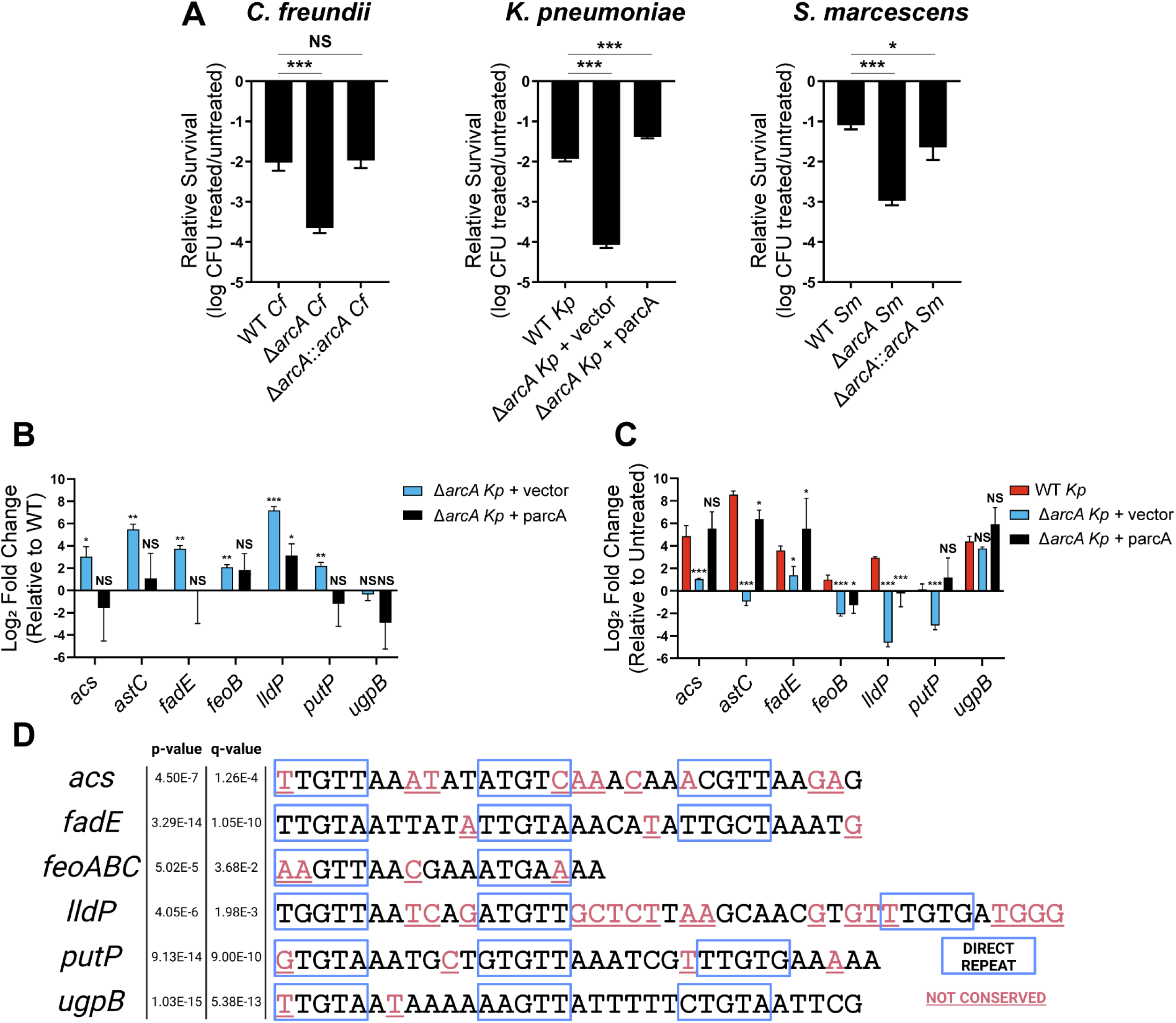
ArcA is involved in the polymyxin B response. **(A)** Overnight cultures grown in LB medium were sub-cultured into LB medium and incubated aerobically to mid-log phase. Cultures were normalized to an OD_600_ 0.2 and treated with polymyxin B for one hour at 37°C. Survival was assessed relative to untreated cultures, and the log transformed data are presented as an average of technical triplicates. Each graph is representative of three independent experiments. Significance was determined by comparing the wild-type strain with the mutant and complemented strains with Dunnett’s multiple comparisons test. **(B-C)** To measure expression of candidate ArcA-regulated genes in the wild-type, *arcA*, and complemented *arcA* KPPR1 strains, mid-log phase cells grown in LB were normalized to approximately 2 x 10^8^ CFU/mL in PBS. Cells were treated with 5µg/mL polymyxin B for 15 minutes followed by RNA extraction. RT-qPCR was performed to assess expression of *acs, astC, fade, feoB, lldP, ugpB* with *gap* serving as the housekeeping gene. Results are displayed as log_2_ fold change and are the average of 3 biological replicates with standard deviation. **(B)** In untreated conditions, expression of each gene by the mutant and complemented strains was compared to that of the wild-type strain following normalizing of Ct values to *gap* and log transformation. Significance was determined via a one-sample *t*-test with a null hypothetical value of zero. **C)** Expression of each gene was then compared between untreated and polymyxin B conditions for each strain. Significance was determined by comparing the wild-type strain with the mutant and complemented strains with Dunnett’s multiple comparisons test. **(D)** FIMO was utilized to search for ArcA binding boxes from *E. coli* K-12 MG1255 (18) in the promoter regions of the seven genes evaluated in the expression studies. Sequences that had a *p-*value and *q-*value at or below 0.05 were considered significant. In the promoters of 6/7 *K. pneumoniae* genes, a putative ArcA binding sequence was identified. Underlined, red nucleotides were loci not conserved between *E. coli* and *K. pneumoniae* sequences. Direct repeats within sequences were labeled based on coordinates of direct repeats within corresponding promoters of *E. coli* genes and are denoted by blue boxes. *p*-values: *≤0.05, **≤0.01, ***≤0.001, NS = not significant

A published transcriptome of *K. pneumoniae* of PMB responsive genes (44) was compared to an established *E. coli* ArcA regulon (18) to identify putative conserved transcripts controlled by ArcA in response to PMB. *acs*, *astC*, *fadE, feoB, lldP, putP,* and *ugpB* were selected for this study based on amino acid identity of at least 80% between *E. coli* CFT073 and *K. pneumoniae* KPPR1. Gene expression was measured by qRT-PCR in mid-log growth for wild-type, *arcA* mutant, and complemented *arcA* mutant of *K. pneumoniae* cells following treatment with a sublethal dose of polymyxin B for 15 minutes. In untreated conditions, every gene except *ugpB* was more highly expressed in the *arcA* mutant relative to the wild-type strain, confirming the ability of ArcA to repress these transcripts (**Fig. 6B**). In all cases, genetic complementation reduced transcript levels compared to the *arcA* mutant. Expression of the same genes was then measured in the presence of polymyxin B. Upregulation of *acs*, *astC*, *fadE, feoB, lldP*, and *ugpB* was observed relative to untreated bacteria in the wild-type cells ranging from 2.0-fold to more than 375-fold **(Fig. 6C)**. In contrast, *putP* exhibited minimal polymyxin B induction. The complemented strain yielded largely similar results to wild-type except for *feoB* and *lldP* in which an intermediate phenotype was noted. Relative expression of *acs* and *fadE* were 30.8 and 3.5 times lower in the *arcA* mutant cells in comparison to wild-type levels yet were still upregulated. Expression levels compared to wildtype were lower for *astC* (3.0 logs), *feoB* (0.9 logs), *lldP* (2.3 logs), and *putP* (1.2 logs) in the *arcA* mutant, and these genes were ultimately downregulated following polymyxin B treatment. In summary, ArcA is largely a repressor of the tested genes in untreated conditions but clearly serves as an activator or mediates de-repression in response to polymyxin B-induced stress.

Given the evidence for ArcA-dependent regulation of the *K. pneumoniae* polymyxin B-induced transcripts, the potential for proximal ArcA binding sites was explored. ArcA binding sequences with two to four direct repeats have been reported for the seven genes of interest in *E. coli* (18). A homologous sequence was identified in six of the seven genes in *K. pneumoniae* (**Fig. 6D**). Coordinates of the direct repeats in the *E. coli* ArcA-binding sequences were then mapped onto the *K. pneumoniae* sequences. The spacing of ArcA binding capabilities was hypothesized remain the same based on the high conservation of ArcA’s structure. Remarkably, most of the nucleotide differences between the *E. coli* and *K. pneumoniae* sequences were outside of the direct repeats, suggesting a pressure for conservation of the direct repeats. Putative ArcA binding sequences were also readily identifiable in many of the same genes of *C. freundii* and *S. marcescens* (**Fig. S3**). The polymyxin B survival assay, expression data, and identification of putative ArcA binding sites in the promoters of polymyxin B-responsive genes all provide evidence for a direct role of ArcA in responding to CAMPs, further emphasizing the function of ArcA in the infection environment.

### ArcA responds to electron transport chain perturbations to promote fermentation

ArcA represses pathways that ultimately provide the electron transport chain (ETC) with electron carriers such as NADH for chemiosmotic-based ATP production (18,49). The ability of the ETC to maintain a proton gradient across the inner membrane can be compromised when the cell envelope is damaged. Thus, ArcA is hypothesized to repress pathways that fuel the ETC in *C. freundii*, *K. pneumoniae*, and *S. marcescens* when proton motive force (PMF) cannot be maintained despite the availability of electron donors and a terminal electron acceptor. The PMF uncoupler carbonylcyanide-*m*-chlorophenylhydrazone (CCCP) was utilized to probe the cell’s ability to respond to inhibition of ATP production via chemiosmosis. Wild-type, *arcA* mutant, and complemented strains were cultured aerobically in a minimal medium containing glucose with and without CCCP to test this hypothesis **(Fig. 7A-B)**. The differences in growth patterns during CCCP treatment varied by species but can be broadly characterized as detrimental. The *arcA* mutants of *C. freundii*, *K. pneumoniae*, and *S. marcescens* had increased lag times of 8.0 h., 4.2 h., and 8.3 h. and 25.2 min., 32.1 min., and 18.7 min. longer doubling times relative to the wild-type strains, respectively.

**Fig. 7:**
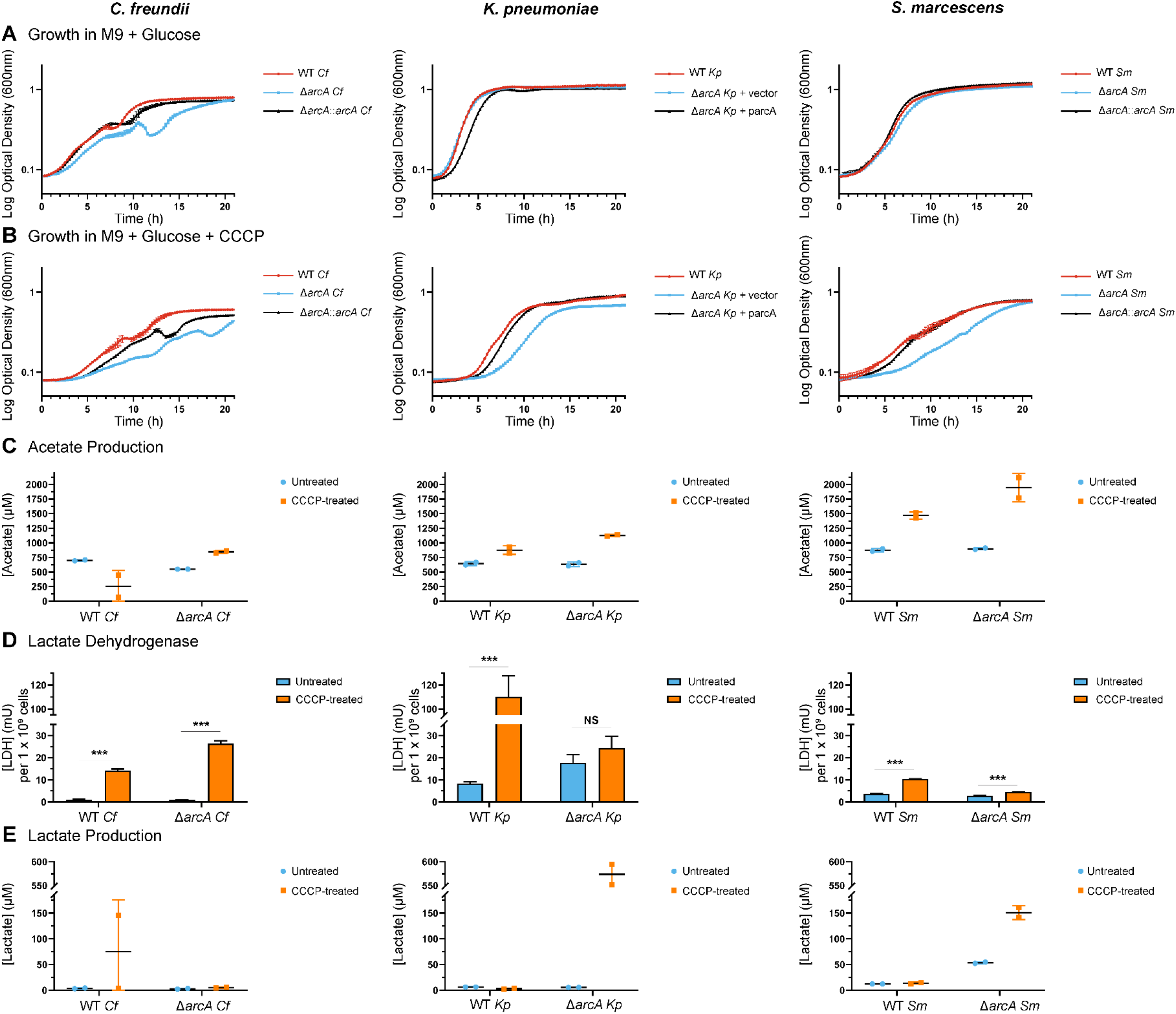
ArcA modulates metabolism in response to disruption of proton motive force by the uncoupler carbonylcyanide-m-chlorophenylhydrazone (CCCP). The ability of wild-type and Δ*arcA* mutant cells to respond to disruption of ATP synthesis via oxidative phosphorylation despite the availability of glucose and oxygen was tested. Overnight cultures incubated aerobically in LB were inoculated into M9 minimal medium with 0.4% glucose without (A) or with (B) CCCP (*C. freundii*, 15µM CCCP; *K. pneumoniae*, 20µM CCCP; *S. marcescens*, 25µM CCCP). Cultures were incubated at 37°C under aerobic conditions and growth was tracked via OD_600_ by a plate reader every 15 minutes. Growth curves are the average of technical triplicates with standard deviation and are representative of three independent experiments. (C) Targeted metabolomics by LC-MS was utilized to quantitate acetate from supernatants of wild-type and *arcA* mutant cultures in early exponential phase from the same conditions as the growth curves. The average of two biological samples with standard deviation are presented in each graph. (D) d-Lactate dehydrogenase (d-LDH) was measured from cell lysates of cultures grown in M9 minimal medium with 0.4% glucose without or with CCCP at the same concentrations as the growth curve conditions. d-LDH levels were quantified with Amplite® Fluorimetric D-Lactate Dehydrogenase Assay Kit (AAT Bioquest) by comparing sample readings to known standards. d-LDH levels were normalized per 1 x 10^9^ cells. The average of three technical replicates with standard deviation is presented as representative of three independent experiments. LDH levels were compared for strains in untreated and treated conditions using Šídák’s multiple comparisons test to determine significance. (E) Targeted metabolomics was repeated to quantify lactate with the same experimental set-up as acetate (C). See methodology for details, Fig. S4 for sampling metrics, and Fig. S5 for LC-MS acetate and lactate samples. *p*-values: *≤0.05, **≤0.01, ***≤0.001, NS = not significant

The ability to grow in CCCP is expected to require an ETC-independent mechanism for ATP production, such as fermentation. ArcA mediates the transition to fermentation (50), so *arcA* mutant bacteria were hypothesized to experience defects in mixed acid fermentative processes in response PMF uncoupling (51–54). LC-MS was utilized to quantify acetate (**Fig. S4**) in the supernatant of untreated and CCCP-treated cultures (**Fig. S5**) as one readout of fermentation. Acetate levels decreased in wild-type *C. freundii* 6.2-fold but were 1.5 times higher in the corresponding *arcA* strain relative to untreated conditions (**Fig. 7C**). In the *K. pneumoniae* and *S. marcescens* wild-type and *arcA* strains, acetate levels were 1.4 to 2.2-fold higher in CCCP-treated conditions (**Fig. 7C**), signifying fermentation was induced in these cultures. D-lactate dehydrogenase levels (LDH) in the CCCP cultures were also measured as an additional metric of fermentation (**Fig. 7D**). LDH activity significantly increased in the *C. freundii* wild-type (13.5-fold) and *arcA* mutant (30.3-fold) strains cultured with CCCP relative to untreated conditions, indicating CCCP induced fermentation, but ArcA activity may play an inhibitory role of LDH in this case. Relative LDH levels also increased in wild-type *K. pneumoniae* (13.2-fold) and *S. marcescens* (2.8-fold) cultures containing CCCP, and importantly, the increase in LDH activity was dependent on ArcA for *K. pneumoniae* and partially so for *S. marcescens*. The relationship between cellular LDH levels and supernatant lactate concentration in the context of CCCP and ArcA was assessed by quantifying lactate by LC-MS. The concentration of lactate increased 20.7-fold in the wild-type *C. freundii* in response to CCCP, but this phenotype was variable and was not observed in the *arcA* mutant strain (**Fig. 7E**). Almost no difference in lactate was found between the wild-type *K. pneumoniae* cultures whereas the *K. pneumoniae arcA* mutant CCCP culture yielded 2 logs more lactate than the untreated culture. A similar trend was observed for *S. marcescens* in which lactate levels did not change between CCCP and untreated wild-type strain cultures but almost tripled for the *arcA* mutant strain (**Fig. 7E**). Lactate levels have previously been shown to increase in the supernatant of *arcA* mutant cultures under anaerobic conditions (55,56), indicating our findings for *K. pneumoniae* and *S. marcescens* matched other fermentative conditions. The inverse correlation of higher LDH levels in wild-type cells to lower lactate concentrations for *K. pneumoniae* and *S. marcescens*, however, are not clear but may potentially be explained by an unknown effect of CCCP treatment or oxidation of lactate at the transport chain (57).

## DISCUSSION

The two-component response regulator ArcA is highly conserved among Enterobacterales species and mediated metabolic adaptation under low oxygen levels in *C. freundii*, *K. pneumoniae*, and *S. marcescens.* We demonstrate for the first time that ArcA promotes fitness of all three species during bacteremia. *arcA* mutants exhibited a dysregulated response to changes in oxygen and iron availability, which are conditions that are likely to be encountered during infection. ArcA was found to be part of the response to membrane damage caused by the CAMP polymyxin B, demonstrating an expanded role for ArcA that is perhaps linked to disruption of ETC activity. ArcA mediated a shift to fermentation in response to PMF disruption, independent of oxygen availability, as measured by LDH activity. The proposed model detailing ArcA’s response to low oxygen, limited iron, and membrane damage is summarized in **Fig. 8**.

**Fig. 8:**
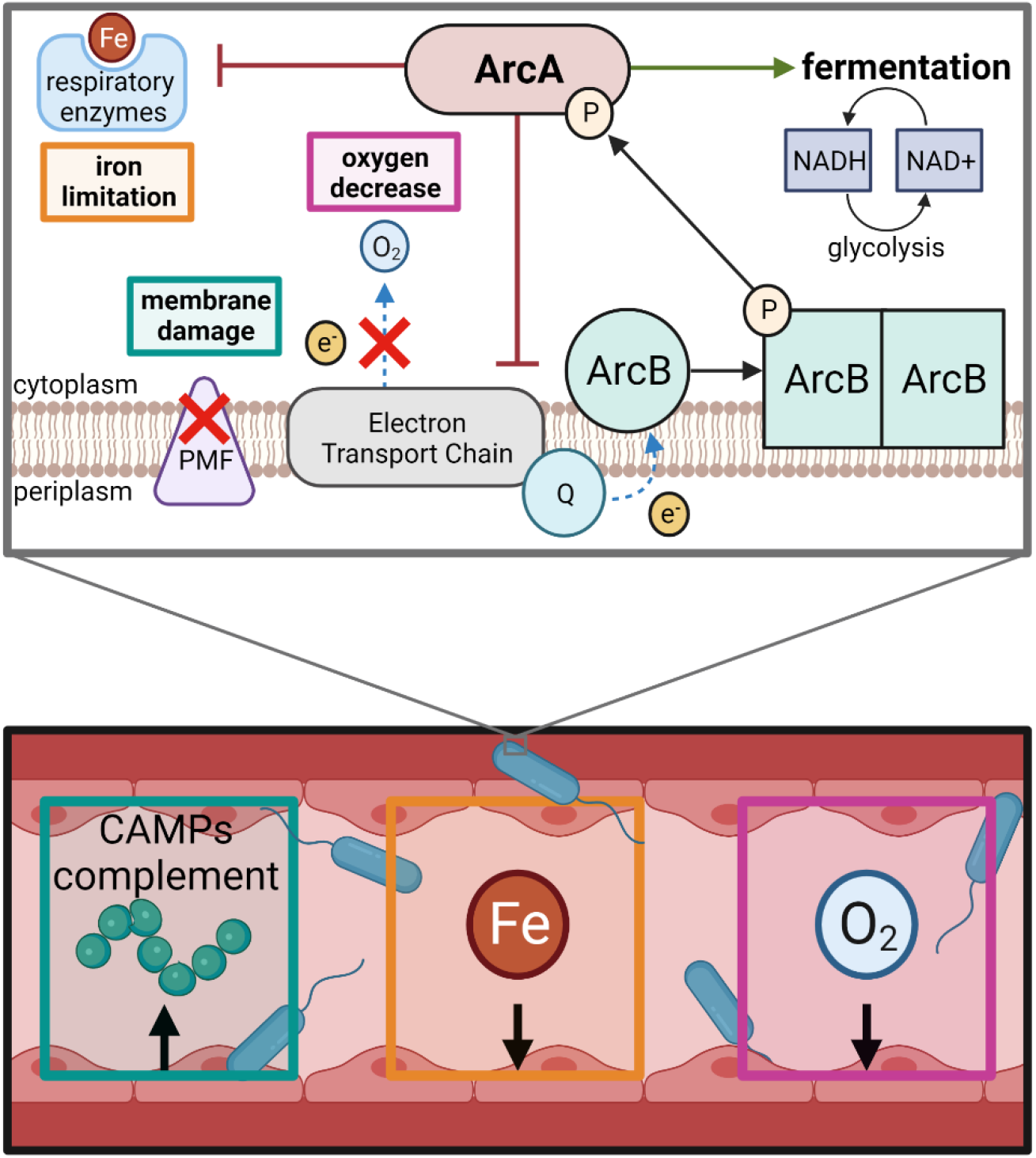
Response regulator ArcA supports fitness during Gram-negative bacteremia. Within the mammalian bloodstream, bacteria encounter decreased iron (Fe) availability, oxygen (O_2_) levels, and elements of the host immune system such as cationic antimicrobial peptides (CAMPs) which can cause membrane damage. ArcA mediates the transition to fermentation in response to such conditions unfavorable for respiration including the inability to maintain a proton motive force (PMF). Quinones (Q) of the electron transport chain transfer electrons to sensor kinase ArcB instead of to pathways which lead to oxygen as the terminal electron acceptor. ArcB then phosphorylates and activates ArcA in response to decreased electron transport chain activity, providing a mechanism by which ArcA can respond to multiple stimuli impacting metabolic activity within the cell.

Bacteria entering the bloodstream from the environment or from another infection site such as the lungs during pneumonia can be hypothesized to experience increasingly anaerobic conditions during dissemination. Ambient oxygen levels are at approximately 21.1%, and the percentage of oxygen in the host decreases to 13.2% in arterial blood to 5.4% in the liver (58). Very little oxygen is dissolved in bloodstream as 98% is bound to hemoglobin (59). A published study from our group has demonstrated the average population doubling time of *C. freundii*, *K. pneumoniae*, and *S. marcescens* in the murine spleen during bacteremia are 66, 39, and 61 minutes, respectively (31). The ability of the bacterial cells to maintain rapid replication rates is thus hypothesized to be an important factor in combating host clearance mechanisms and establishing infection during bacteremia. It is notable that the *in vitro* growth defects observed in this study for the *K. pneumoniae* and *S. marcescens arcA* mutants were evident by a sizeable shift in growth curves in the aerobic to anaerobic transition. These results capture ArcA’s role in responding to a change in oxygen utilization and showcase ArcA’s likely support of the metabolism needed in the host environment to maintain rapid growth. Of note, our research group has previously shown that during urinary tract infections, *E. coli* relies on the TCA cycle with glycolysis being dispensable (60,61). If *E. coli* favors the same pathways during bacteremia, ArcA would be expendable in its role as a repressor of the TCA cycle, which explains the lack of fitness defect associated with mutating *arcA* in the *E. coli* bacteremia model. The requirement of ArcA for the other species suggests other metabolic pathways are likely preferred during infection. Indeed, our previous TnSeq screens identified genes encoding 6-phosphofructokinase, phosphate acetyltransferase, and acetate kinase as contributing to fitness for *C. freundii* and *S. marcescens*, suggesting glycolysis and fermentation are required by these species during infection (10,11).

This work establishes that ArcA is needed to maximize replication of *C. freundii*, *K. pneumoniae*, and *S. marcescens* in iron-limited conditions. ArcA has previously been shown to contribute to iron homeostasis in conjunction with FNR and Fur in *E. coli* (62). The decrease in growth *arcA* mutants observed here occurred under ambient oxygen conditions, bolstering the notion that ArcA responds to utilization of oxygen as a terminal electron acceptor rather than the absence of oxygen. Fermentation has indeed already been shown to be the preferred metabolic pathway during iron starvation (37,63). Chareyre *et al.* demonstrated that when facing iron starvation, *E. coli* shuts down respiratory complexes via the small RNA RhyB. RhyB post-transcriptionally controls the *nuo* and *sdh* operons which encode these complexes. The *nuo* and *shd* operons are well established as being among the genetic elements most strongly repressed by ArcA (18,19). Future studies may examine the potential for coordination between RhyB and ArcA to further our understanding of repression of respiration in the context of iron limitation. This link between iron and oxygen availability appears to be intrinsic to life. Hypoxia Inducible Factor (HIF) in humans is a transcriptional activator induced by low levels of systemic oxygen availability (64). Similarly to ArcA, HIF has also been shown to become active in low iron conditions and promote glycolytic activity (65,66).

Increased sensitivity *C. freundii* and S*. marcescens arcA* mutants to human serum and a lack of growth for a*rcA K. pneumoniae* in heat-inactivated human serum are indications ArcA mediates survival in the bloodstream environment. The studies here are the first to our knowledge to demonstrate bacteria lacking *arcA* are more sensitive to CAMPs. Six genes upregulated by ArcA were identified in *K. pneumoniae* following PMB treatment, which was unexpected given ArcA’s well established role in repressing all of the genes except *feoB* (18–20). There is precedence, however, for the ability of ArcA to upregulate and downregulate the same gene(s) depending on growth conditions (67). More work is needed to determine if ArcA directly or indirectly upregulates the genes responsive to PMB. Interestingly, *acs*, *lldP*, and *putP* are upregulated in an *arcA* avian pathogenic *E. coli* strain grown in duck serum relative to the isogenic wild-type strain where membrane stressors are also likely present (68). None of the seven genes studied in the context of PMB are directly involved with major systems of aerobic respiration. A “core” ArcA regulon may exist in which ArcA repression of central carbon metabolic pathways is invariable alongside a “conditional” regulon in which ArcA’s role as an activator or repressor is context dependent. More transcriptomic and DNA footprinting studies will be critical for defining the direct and indirect ArcA regulons for more species in infection-relevant conditions.

CAMPs such as PMB can damage the inner membrane and inhibits respiratory enzymes (69,70), implying PMB can disrupt maintenance of PMF or damage the ETC itself. In our studies, CCCP was used to directly target ETC function. *arcA* mutants grew more slowly in the presence of CCCP, connecting ArcA to disruptions of ETC activity. CCCP induced an increase in LDH levels in all three species, which was indicative of a shift to fermentation, and this increase was at least partially ArcA-dependent for *K. pneumoniae* and *S. marcescens*. Targeted metabolomics revealed lactate and acetate levels in a medium with CCCP are ArcA-and species-dependent. Acetate and lactate pathways contribute to the maintenance redox balance during glycolysis (54). Based on ArcA’s multiple roles in also maintaining intracellular redox balance (14), acetate and lactate production may be reflective of redox levels in the presence of CCCP. Cells that are more efficient in energy production and carbon cycling may reuse end products of fermentation rather than secrete them into the supernatant. Decreased metabolic efficiency was found in an *arcA* mutant of *E. coli* undergoing anaerobic fermentation based on a 15.8% lower growth rate relative to the wild-type strain (19). *K. pneumoniae* and *S. marcescens arcA* mutants excreted higher levels of acetate in CCCP, which does differ from *E. coli arcA* mutants in other studies which produced the same level of or less acetate than wild-type strains during anaerobic fermentation (55,56,71). Disruption of PMF by CCCP may thus induce fermentative conditions differently than anaerobiosis, or ArcA-mediated fermentation is species specific.

We conclude that ArcA responds to low oxygen conditions, decreased iron levels, and host-mediated membrane damage during bacteremia in three Gram-negative bacterial pathogenic species. Activation of ArcA in response to low iron and membrane damage was not tested, so it remains to be determined if ArcA function is controlled in these contexts by conventional models. ArcA has recently been shown to become partially active independently of ArcB via intramolecular disulfide bonding in oxidizing conditions, providing evidence of additional regulatory mechanisms to be explored (72). Further studies of ArcA in the bloodstream environment will be important in understanding the complex regulation of central carbon pathways utilized during pathogenesis and may reveal more shared or unique metabolic capabilities employed by multiple bacterial species during infection.

## METHODS

### Bacterial strains and culture conditions

The bacterial strains utilized in this study are listed in **Table 1**. *E. coli* TOP10 cells were used for routine cloning purposes. Overnight culture was performed in LB (73) and experimental cultures were grown in LB or M9 medium (74) containing 100µM CaCl_2_, 1mM MgSO_4_, 0.4% D-glucose, and 0.1% casamino acids as indicated. All cultures were maintained at 37°C with 200RPM shaking unless noted otherwise. All anaerobic cultures were maintained in a 37°C anaerobic chamber maintained at 10% H_2_, 5% CO_2_ and 85% N_2_.

### Strain engineering

*C*. *freundii, E. coli, K. pneumoniae,* and *S. marcescens arcA* mutants were generated using Lambda red mutagenesis as previously described (10,75,76) using the oligonucleotides listed in **Table S1.** Chromosomal mutations were confirmed by PCR-amplification and sequencing of the mutant allele. *C. freundii* UMH14 and *S. marcescens* UMH9 Δ*arcA::npII* mutant alleles were complemented by re-integration of the wild-type allele via recombineering. Primers were designed to amplify the portion of the *arcA* gene replaced by the antibiotic resistance cassette in the Δ*arcA::npII* mutants with the same homologous ends via PCR with Q5 polymerase (New England Biosciences). *C. freundii* and *S. marcescens arcA* mutant strains were transformed with the *arcA*-containing PCR products. Recovery of cells was performed in LB without selection at 30°C. Transformants were subsequently passaged in LB multiple times to enrich for complemented mutants via restoration of wild-type growth rates. Enriched populations were then plated onto LB agar and candidate colonies were scored for reversion to the wild-type colony size. Complementation was confirmed via Sanger sequencing of PCR products amplified from the *arcA* locus. The *K. pneumoniae* KPPR1 *arcA* mutant strain was complemented *in trans* using the pBBR1MCS-5 broad host-range plasmid (77). Primers ANB21F and ANB21R and were used to amplify the *arcA* ORF and 539 base pairs upstream of the start of the gene with Easy A polymerase (Agilent). The PCR product and pBBR1MCS-5 parent plasmid were separately digested with SacI and XbaI. Ligation of the two digested fragments was achieved with T4 DNA ligase (NEB) followed by electroporation into *E. coli* TOP10 (Thermo Fischer). Plasmid construction was confirmed by Sanger sequencing. KPPR1 *ΔarcA::npII* was transformed with complementation plasmid by electroporation the complementation or empty vector control plasmids were maintained in the presence of gentamicin (10 µg/ml).

### Murine bacteremia model

Overnight LB cultures of wild-type and a*rcA* mutant constructs were sub-cultured into fresh LB and cultured at 37°C with 200 RPM shaking until mid-log phase. Mid-log cultures were washed and resuspended with PBS and normalized based on OD_600_ to achieve the approximate concentration of bacteria: *C. freundii*: 1 x 10^9^ CFU/mL*, E. coli*: 2 x 10^7^ CFU/mL, *K. pneumoniae*: 1 x 10^6^ CFU/mL, and *S. marcescens*: 1 x 10^8^ CFU/mL. Wild-type and *arcA* mutants for each species were mixed 1:1, and 6 to 8 weeks old male and female C57BL/6 mice (Jackson Laboratory, Bar Harbor, ME) were inoculated via tail-vein injection as previously described (78). Inocula and organ homogenates were plated on LB agar and LB agar containing kanamycin (50 µg/ml) for differential CFU determinations. A competitive index was calculated for each organ by dividing the ratio of mutant to wild-type CFU in the organ by the ratio of mutant to wild-type CFU in the inoculum. Competitive indices were log-transformed, and significance was determined by a one-sample *t*-test with a hypothetical value of zero. Murine experiments were performed in compliance with an animal protocol (PRO00010856) approved by the University of Michigan Institutional Animal Care & Use Committee.

### In vitro growth

Aerobic and anaerobic overnight cultures were normalized based on OD_600_, washed and resuspended with PBS, and subcultured 1:100 into the desired media. For aerobic growth studies, 300µL from each prepared culture was added in triplicate to a honeycomb plate. For iron-limitation cultures, iron-limited M9 media with and without iron supplementation were prepared for each species individually: *C. freundii* – 0.6mM 2,2’-dipyridyl (0.6% DMSO) and 3.0mM FeSO_4_; *K. pneumoniae* – 0.2mM 2,2’-dipyridyl (0.2% DMSO) and 0.1mM FeSO_4_; *S. marcescens* – 0.4mM 2,2’-dipyridyl (0.4% DMSO) and 0.2mM FeSO_4_. Growth was assessed by comparing area under the curve (AUC) in each condition of the mutant and complemented strains to the AUC of the wild-type strains, and significance was determined with Dunnett’s multiple comparisons test. For cultures containing carbonylcyanide-*m*-chlorophenylhydrazone (CCCP), M9 media including the following concentrations were prepared for each species: *C. freundii* - 15µM CCCP, *K. pneumoniae* - 20µM CCCP, *S. marcescens* - 25µM CCCP. Plates were incubated on a Bioscreen-C plate reader with the following settings: 37°C, intermediate continuous shaking, OD_600_ measurement every 15 minutes. For anaerobic growth studies, 200µL from each prepared culture was added in triplicate to a 96 well plate. The plate was incubated in an anaerobic plate reader (BioTek Powerwave HT) with the following settings: 37°C, no shaking, OD_600_ measurement taken every 10 minutes.

### Survival assays

Pooled human complement serum (Innovative Research Lot #37600) was stored at -80C and thawed directly prior to use and heat-inactivated at 56°C for 45 minutes, where indicated. Bacteria cultured to mid-log phase in LB medium were washed and resuspended to a final density of 2x10^8^ CFU/ml in PBS then added to serum in triplicate in 96-well plates. Serum sensitivity was tested a concentration of 10% (*C. freundii*), 90% (*K. pneumoniae*), or 20% (*S. marcescens*) at 37°C. Bacterial viability was determined after a 90-min exposure by serial dilution and colony counts relative to time zero. For polymyxin B studies, cells were collected by centrifugation and resuspended in PBS to an OD_600_ of 0.2. Polymyxin B (RPI Lot# 85594-90055) was added to cells triplicate96-well plates at a final concentration of 5.0µg/mL (*C. freundii*), 50µg/mL (*K. pneumoniae*), or 100µg/mL (*S. marcescens*) with water serving as the control. Plates were stored statically for 1 hour at 37°C followed by enumeration of viable bacteria relative to time zero. For both assays, Dunnett’s multiple comparisons test was used to assess statistical significance following log transformation of data.

### Gene expression

Mid-exponential phase aerobic bacteria were normalized to 2x10^8^ CFU/mL in PBS. 10mL of each resuspended culture was added to a 125mL flask, and 1.0mL of resuspended culture was kept as an untreated control. Polymyxin B (50uL) was added to each flask for a final concentration of 5.0µg/mL. Flasks were then incubated at 37°C and 200 RPM shaking for 15 minutes. 1.0mL of treated and untreated culture were added directly to 2mL of RNA protect solution (Qiagen), and RNA was extracted with the RNeasy Mini Kit (Qiagen) following manufacturer’s instructions. RNA samples were treated with RQ1 DNase (Promega) and repurified with the RNeasy Mini Kit (Qiagen). cDNA was generated with iScript cDNA Synthesis Kit (Bio-Rad) and diluted 1:10 with water. RT-qPCR was performed with Power SYBR Green (Thermo Fischer) followed by calculation of relative gene expression with the 2^-ΔΔCt^ (Livak) method (79). In untreated conditions, gene expression was compared to the wild-type strain following log transformation, and significance was determined via a one-sample *t*-test with a null hypothetical value of zero. Expression of each gene was compared between untreated and polymyxin B conditions, and significance was determined by comparing the wild-type strain with the mutant and complemented strains with Dunnett’s multiple comparisons test.

### Metabolite quantification

The experimental set-up was the same as that of the CCCP growth curve studies with the addition of two wells per strain and condition for sampling. OD_600_ readings were monitored in real time via the Bioscreen-C plate reader. Upon reaching early exponential phase, 300uL was removed from two wells for each strain and condition and immediately transferred to ice. An aliquot of each sample was removed for CFU enumeration before cells were pelleted in a 4°C microcentrifuge. The supernatant was transferred to a new tube and immediately stored at -80°C. Supernatant samples were processed by the University of Michigan Metabolomics Core to quantify acetate and lactate.

Short chain fatty acids (SCFAs), including acetate, were measured using a modified version of a previously described protocol (80). SCFAs in the sample supernatant were derivatized using 3-nitrophenylhydrazine and an EDAC-6% pyridine solution. Samples were analyzed via LC-MS alongside acetate controls ranging from 3µ to 3000µM using an Agilent (Santa Clara, CA) 1290 LC coupled to an Agilent 6490 triple quadrupole MS. The chromatographic column was a Waters (Milford, MA) HSS T3, 2.1 mm x 100 mm, 1.7 µm particle size. Quantitation was performed using Agilent MassHunter Quantitative Analysis software version 8.0 by measuring the ratio of peak area of the 3-NPH derivatized SCFA species to its closest internal standard.

Lactate quantification was performed starting with the addition of an extraction solvent containing ^13^C Lactate to each supernatant sample. Following a series of mixing and centrifugation, supernatant was collected and dried using a nitrogen blower. Samples were reconstituted alongside a series of calibration standards. Ion pairing reverse phase LC-MS analysis was then performed using an Infinity Lab II UPLC coupled with a 6545 QTof mass spectrometer (Agilent Technologies, Santa Clara, CA) and a JetStream ESI source in negative mode. Chromatographic separation was performed on an Agilent ZORBAX RRHD Extend 80Å C18, 2.1 × 150 mm, 1.8 μm column with an Agilent ZORBAX SB-C8, 2.1 mm × 30 mm, 3.5 μm guard column. Data were processed using MassHunter Quantitative analysis version B.07.00.

### Lactate dehydrogenase measurement

Bacteria were cultured as described for metabolite quantification and early exponential phase cells were collected via centrifugation at 4°C. The supernatant was removed, and cells were washed once then resuspended in PBS at 4°C prior to sonication. Cells were lysed by sonication with a taper microtip Z192740-1EA (Sigma-Aldrich) with the following protocol: 1 min. 40s. sonication at 40% amplitude with 4 s. bursts divided by 10 s. pauses to avoid overheating. Lactate dehydrogenase was measured in triplicate from cleared lysates with the Amplite® Fluorimetric D-Lactate Dehydrogenase (LDH) Assay Kit from AAT Bioquest per the manufacturer’s instructions. Fluorescence (excitation: 540nm; emission: 590nM) was measured after one hour incubation at room temperature protected from light with a Synergy H1 plate reader. LDH was quantified in samples based on standards ranging from 1µM/mL LDH to 200µM/mL. LDH concentration per 1 x 10^9^ cells was subsequently normalized based on CFU enumeration and compared for each strain between untreated and CCCP-treated conditions. Significance was determined by comparing LDH levels for strains in untreated and treated conditions using Šídák’s multiple comparisons test.

### In silico analyses

ArcA amino acids sequences (n=419) from 418 Enterobacterales species were collated from BV-BRC (81) (**File S1**). A multi-sequence alignment was generated with MUSCLE via EMBL-EBI (82,83). An ArcA predicted structure AF-P0A9Q1-F1 from Alpha Fold was retrieved via UniProt to serve as a template for conservation mapping (23,24,84). Consurf calculated conservation scores from the multiple sequence alignment based on the sequence extracted from the predicted structure via a JTT evolutionary model (**File S2**) (85,86). Conservation scores were mapped onto the predicted ArcA structure by Consurf with visualization of this projection provided by PyMOL (26). The ArcA amino acid sequences of *C. freundii* UMH14, *E.* coli CFT073, K*. pneumoniae* KPPR1, and *S. marcescens* UMH9 were aligned with Clustal Omega (30). A percent identity matrix was generated to calculate pairwise percent identities for all possible combinations of the four species. The output of amino acid alignment between all four species was then examined to assess conservation. Similarity of non-conserved residues was defined according to set parameters with a Gonnet PAM 250 matrix score of >0.5 signifying “strongly similar” and a score <0.5 and >0 for “weakly similar” residues.

ArcA binding sequences in the promoters of *acs*, *astC*, *fadE*, *feoB*, *lldP*, *putP*, and *ugpB* from *E. coli* K-12 MG1655 were used as the input motif with which to scan the promoters of the same genes in *C. freundii*, *K. pneumoniae*, and *S. marcescens* (18). Sequences identified by FIMO Version 5.5.1 from MEME Suite were reported as potential ArcA binding sequences when *p-* values and *q-*values (false discovery rate) were both ≤0.05 (87). Nucleotides of the *E. coli* sequence and the sequences of the other species were compared to assess homology. Putative direct repeats were mapped onto the proposed ArcA sequences based on the coordinates reported in the *E. coli* sequences.

## ACKNOWLEDGEMENTS

We thank current and past members of the Mobley and Bachman laboratories for their continual feedback over the course of these studies. We also thank Dr. Maria Sandkvist for her expertise and insight while planning and interpreting many of the experiments as well as the laboratories of Dr. Nicole Koropatkin and Dr. Eric Martens for technical support and equipment used during the anaerobic studies. The Biomedical Research Core Facilities at the University of Michigan and associated vendors are acknowledged for their services including DNA sequencing and metabolomics. The summary figure was created with BioRender.com. We lastly acknowledge the use of C57BL/6 mice sacrificed during the bacteremia model.

## SUPPORTING INFORMATION

File S1: Table of Enterobacterales genomes and ArcA ORFs analyzed for conservation modeling

File S2: Specific amino acid residue information of ArcA conservation model

Figures S1 – S5 and Table S1 are provided in “Supporting Figures and Table.”

